# High-Throughput Bioprinting of the Nasal Epithelium using Patient-derived Nasal Epithelial Cells

**DOI:** 10.1101/2023.03.29.534723

**Authors:** I. Deniz Derman, Miji Yeo, Diana Cadena Castaneda, Megan Callender, Mian Horvath, Zengshuo Mo, Ruoyun Xiong, Elizabeth Fleming, Phylip Chen, Mark E. Peeples, Karolina Palucka, Julia Oh, Ibrahim T. Ozbolat

## Abstract

Human nasal epithelial cells (hNECs) are an essential cell source for the reconstruction of the respiratory pseudostratified columnar epithelium composed of multiple cell types in the context of infection studies and disease modeling. Hitherto, manual seeding has been the dominant method for creating nasal epithelial tissue models. However, the manual approach is slow, low-throughput and has limitations in terms of achieving the intricate 3D structure of the natural nasal epithelium in a uniform manner. 3D Bioprinting has been utilized to reconstruct various epithelial tissue models, such as cutaneous, intestinal, alveolar, and bronchial epithelium, but there has been no attempt to use of 3D bioprinting technologies for reconstruction of the nasal epithelium. In this study, for the first time, we demonstrate the reconstruction of the nasal epithelium with the use of primary hNECs deposited on Transwell inserts via droplet-based bioprinting (DBB), which enabled high-throughput fabrication of the nasal epithelium in Transwell inserts of 24-well plates. DBB of nasal progenitor cells ranging from one-tenth to one-half of the cell seeding density employed during the conventional cell seeding approach enabled a high degree of differentiation with the presence of cilia and tight-junctions over a 4-week air-liquid interface culture. Single cell RNA sequencing of these cultures identified five major epithelial cells populations, including basal, suprabasal, goblet, club, and ciliated cells. These cultures recapitulated the pseudostratified columnar epithelial architecture present in the native nasal epithelium and were permissive to respiratory virus infection. These results denote the potential of 3D bioprinting for high-throughput fabrication of nasal epithelial tissue models not only for infection studies but also for other purposes such as disease modeling, immunological studies, and drug screening.

## 1. Introduction

The nasal epithelium is composed primarily of pseudostratified columnar epithelial cells containing basal, ciliary, non-ciliary mucus-secreting goblet cells, which play a crucial role in the physiological and immunological functions of the nasal cavity [1]. These cells are the first barrier against substances that are inhaled in daily life, such as allergens, viral, bacterial and fungal pathogens [1–3], and are responsible for [1–3] mucociliary clearance led by a wave action removing foreign bodies and preserving the homeostasis of the nasal cavity [4,5]. Tight junctions between nasal epithelial cells play a crucial role in maintaining the integrity of the nasal epithelial barrier by sealing the spaces between adjacent epithelial cells and preventing the passage of molecules and pathogens through the paracellular space [6,7]. These junctions are made up of transmembrane proteins such as claudins, occludins, and junctional adhesion molecules that interact with each other and with the cytoskeleton to form a stable barrier. The expression and localization of these proteins can be altered by different physiological and pathological stimuli, such as inflammation and allergens. This can lead to changes in barrier function, increased permeability of the epithelium, and the development of inflammatory conditions [7–9].

The use of bronchial epithelial cells obtained from brush biopsies has been a common established method for fabrication of respiratory epithelial models in the literature [10] [10]; however, these cells may not be representative of healthy epithelium due to surrounding inflammation, disease, or possible deformation in the three-dimensional (3D) epithelial structure [11,12]. An alternative source of epithelial cells is from the nasal cavity, which can be obtained through less invasive procedures. The use of freshly isolated primary airway cells or commercially available fully-differentiated respiratory epithelial cell cultures can be costly and limit the variety of donors available [13]. Therefore, in-vitro culture of functional respiratory epithelium is crucial in current respiratory tissue engineering approaches to overcome these limitations [14]. Recently, respiratory epithelial in-vitro systems have become an alternative to expensive, labor-intensive animal studies, which also raise ethical issues, regardless of their relevance to human physiology [15,16]. These in-vitro systems utilize a wide range of cell sources and cultivation protocols allowing the investigation of respiratory diseases and the effects of various environmental factors on the airways [17,18]. Although immortalized cell lines have the potential to overcome limitations of primary cell cultures, they may lack complete differentiation and the ability to generate ciliated cells. Papazian et al. argued that in-vitro differentiation of primary human respiratory epithelial cells may be the most suitable option for studying the airway epithelium despite of its complex and time-consuming process [19,20]. To address such issues, it is important to standardize and validate methods (i.e., protocols, cells, scaffolds, and media) with inter-laboratory reproducibility, ultimately, for replacing animal testing.

Bioprinting is a rapidly growing field that uses the basics of 3D printing technologies to create living structures composed of cells and biomaterials [21–23]. One of the major purposes of bioprinting is to create functional tissue models that can be used for research, drug discovery, and eventually, therapeutic applications. Thus, bioprinting has been used to produce various types of epithelial tissue models such as skin [24], intestinal [25,26], bronchial epithelium [27] and more recently, alveolar epithelium [28,29]. Nevertheless, to date, researchers conventionally use manual cell seeding to develop differentiated cultures of the nasal epithelium [30]. In addition, various other techniques, such as microfluidic channels with a bioreactor [30], spheroid [31,32] and hanging drop culture [33,34], have been used to reconstitute the nasal epithelium composed of multiple cell types. However, manual cell seeding is a slow process, lacks throughput and does not yield uniform tight junctions, and other abovementioned approaches still have a critical limitation to facilitate the development of an anatomically accurate 3D model, in which 3D bioprinting can overcome such limitations. Bioprinting can also facilitate homogeneous cell distribution with high accuracy in a high-throughput manner for investigating nasal disorders and diseases including but not limited to sinusitis, asthma, and viral infection. To the best of our knowledge, bioprinting has not been reported to reconstruct the nasal epithelium in the literature.

In this study, we present the first-time demonstration of nasal epithelial reconstitution via droplet-based bioprinting (DBB) of primary human nasal epithelial progenitor cells (hNECs). To elaborate, DBB was employed by depositing hNEC-laden solution onto Transwell inserts in 24-well plates, in which each layer was deposited in a ‘pass’ over the surface, and the number of passes were varied from 10 to 50 to modulate the cell density per insert. Upon confirmation of the biocompatibility, hNECs were maintained in submerged culture for 10 days to reach confluency and form tight junctions and then at the air-liquid interface for three weeks to fully differentiate. Compared to manually seeded hNECs, bioprinted hNECs, at an even relatively lower cell density, exhibited successful differentiation into multiple cell types, formation of pseudostratified columnar epithelial architecture, and evidence of functionality (i.e., barrier function, mucus secretion, and beating cilia). Overall, the presented technique facilitated the effective fabrication of the nasal epithelial tissue in a high-throughput manner with structural and physiological characteristics like the native nasal epithelium.

## 2. Materials and methods

### 2.1 Harvesting of hNECs

hNECs were collected from the nasal turbinates of consenting volunteers with a brush (CytoSoft CYB-01) as approved by the Institutional Review Board (IRB) at Nationwide Children’s Hospital (IRB numbers 16-00827 and 17–00594). This work was classified as a level 1 risk clinical study—no greater than minimal risk (pursuant under 45 Code of Federal Regulations [CFR] 46.404 and 21 CFR 50.51). Informed consent procedures followed in compliance with Nationwide Children’s Hospital Research Responsible Conduct Guidelines.

### 2.2 Manual initiation of nasal epithelial cultures

Progenitor hNECs were placed in phosphate buffered saline (PBS) with Ca^2+^/Mg^2+^ and stored at 4 °C until processing. They were removed from brushes and treated with a declumping solution containing PBS, dithiothreitol (DTT; 3mM), ethylenediaminetetraacetic acid (EDTA; 2mM), and Type II collagenase (12.5 mg/100 ml) for 5-10 min at 37 °C followed by the addition of 10% fetal bovine serum (FBS). Cells were plated on tissue culture dishes precoated with Type I bovine collagen (PureCol). They were expanded in bronchial epithelial cell growth media (BEGM) [35] or Ex Plus (Stem Cell Technologies) medium for 1 week before being cryopreserved. Upon thawing, the cells were plated on Type IV collagen (Sigma-Aldrich) coated Transwell inserts (Corning 3470; polyester, 6.5 mm diameter, 0.4 µm pores) and used as a control group.

### 2.3 Droplet-based bioprinting (DBB) of the nasal epithelium

For a sterile environment, a customized droplet-based bioprinter (jetlab® 4, MicroFab Technologies Inc.) was placed in a biosafety cabinet (Air Science Purair VLF36) with vertical laminar flow. A micro-valve dispensing device (The Lee Company, cat. no. INKX0517500A) connected with a removable nozzle (250 mm inner diameter, The Lee Compan2y, cat. no. INZA3100914K) was used to expel solutions placed in fluid reservoirs. Then, 10 mg/mLType IV collagen solution was printed for coating the surface of 6.5 mm Transwell inserts in 24-well plates. Here, the printing parameters were set as 5 V dwell voltage, 500 ms dwell time, 1 ms rise/fall time, 0 V echo voltage, 20 ms echo time, 100 Hz frequency, and 126 kPa back pressure. After air-drying the collagen-coated inserts in the biosafety cabinet overnight, hNEC-laden PneumaCult-ALI (Stemcell Technologies) medium (a density of 0.022 × 10^5^ cells/ml) was bioprinted in a 7 mm circular shape layer by layer, in which a single layer was referred to as a “pass” as shown in **Figure 1A**. The hNEC-laden structures were bioprinted in 10, 20, 30, 40, and 50 passes to compare with manually seeded hNECs to explore the role of cell seeding density on the differentiation of hNECs and formation of the nasal epithelium. During our experiments, approximately 2.2 × 10^5^ cells were seeded per insert for the manual seeding group while 0.22, 0.44, 0.66, 0.88 and 1.1 × 10^5^ were bioprinted per insert for the 10-, 20-, 30-, 40- and 50-pass group, which took ∼2, 4, 6, 8 and 10 min, respectively (**Table 1**, **Video S1**).

**Table 1.**
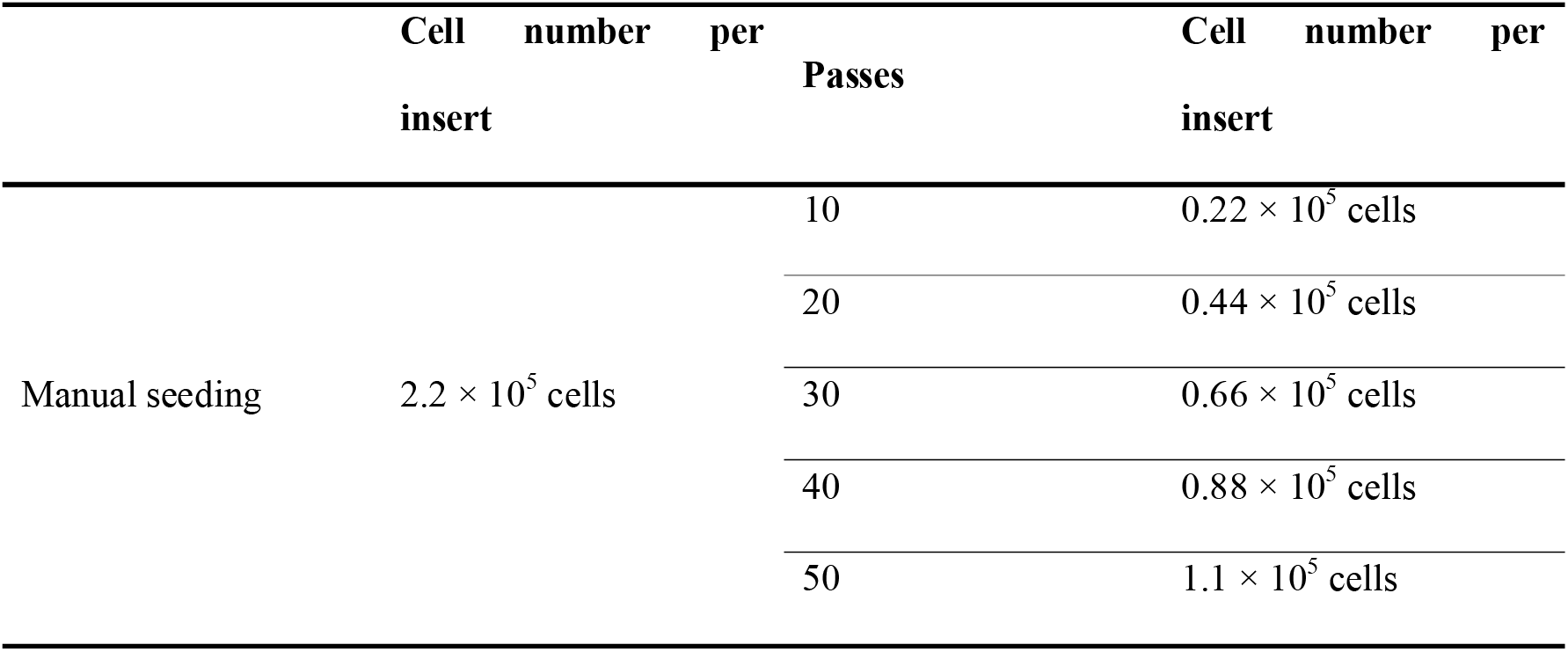
Cell numbers used for manual seeding and DBB.

**Figure 1.**
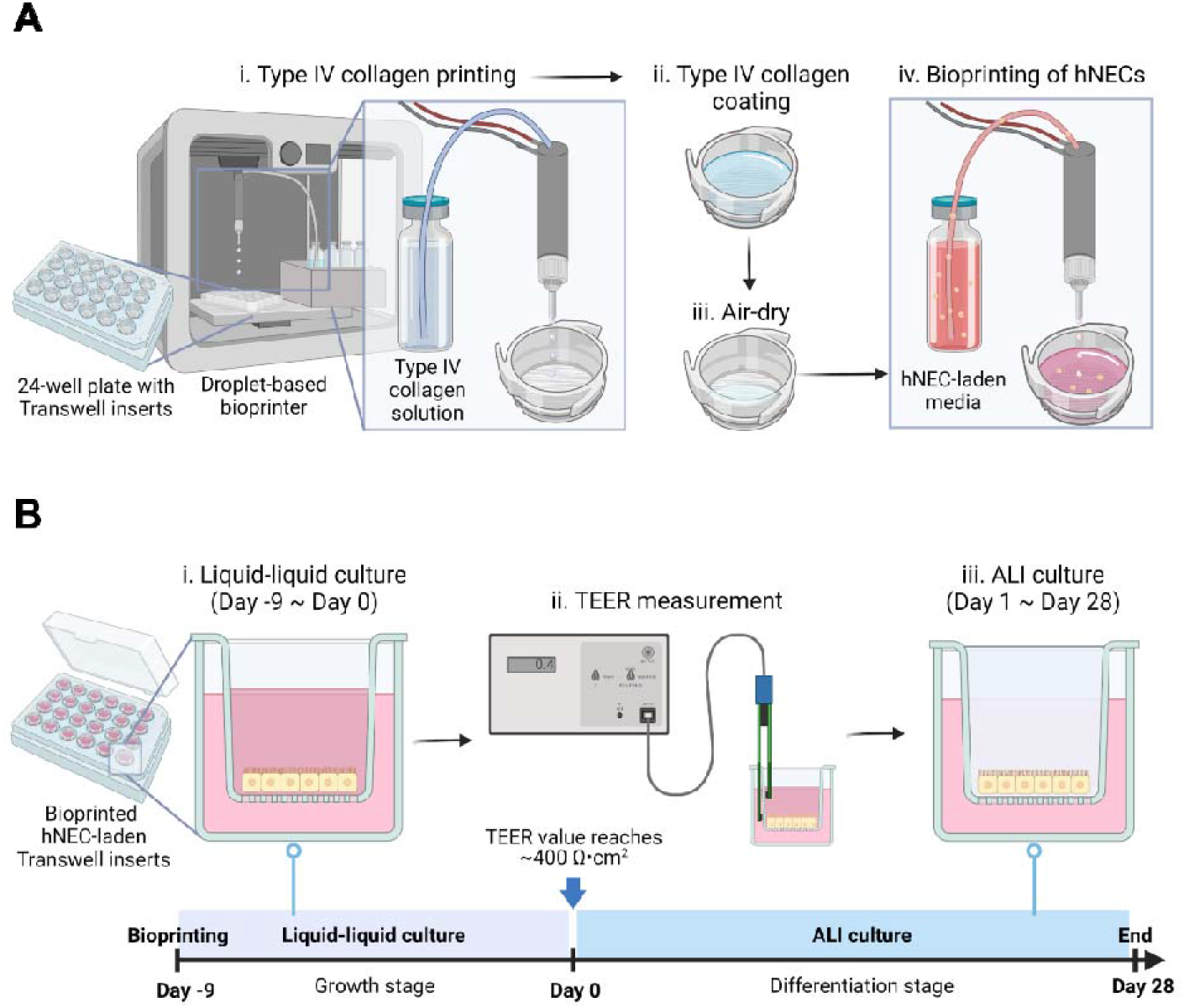
A schematic image illustrating **(A)** DBB of the nasal tissue model and **(B)** its culture along with the timeline for liquid-liquid followed by ALI culture. This figure was created using BioRender (https://biorender.com).

### 2.4 ALI culture of hNECs

Bioprinted and manually prepared Transwell inserts on 24-well plates were cultured for 37 days as shown in **Figure 1B**. The media consisted of PneumaCult-ALI basal medium (Stemcell Technologies) supplemented with PneumaCult-ALI 10×supplement, PneumaCult-ALI Maintenance Supplement (100x), Hydrocortisone stock solution, Heparin solution with the Rock Inhibitor (all from Stemcell Technologies) were mixed and added to both the apical (0.2 mL) and basal (0.65 mL) compartments. The samples were submerged for the first ten days, which were referred as Day -9 to 0 (**Figure 1B**). On Day 0, the apical culture medium was removed from the ALI culture and the basal chamber medium was replaced with the same PneumaCult ALI media but lacking the Rock Inhibitor. This medium was replaced every 2 days until Day 28. Excess mucus was removed from the apical surface by washing the cells once with Dulbecco’s PBS (DPBS 1X; Corning) as required but at least once per week to prevent excessive mucus accumulation.

### 2.5 Cell viability analysis

After DBB, cell viability was measured using LIVE/DEAD staining (Invitrogen, CAT: R37601) before cell differentiation at a day after DBB (Day -8) and a week later (Day -2). A staining solution was prepared by mixing calcein AM (0.15 mM) and ethidiumhomodimer-1 (2 mM) and added to the apical surface of each culture, which was placed in the incubator with 5% CO_2_ at 37 ^°^C for 30 min. The stained cells were imaged using an Axio Zoom fluorescent microscope (Zeiss). The percent (%) cell viability was calculated by dividing the number of live cells by the total number of cells and multiplying by 100.

### 2.6 Immunofluorescent staining

The variations in Zonula Occludens-1 (ZO-1), cilia expression and mucus positivity were examined using immunofluorescent imaging. hNECs were being fixed with 4% paraformaldehyde (PFA) overnight at 4 ^°^C. They were permeabilized with 0.1% Triton X-100 (Sigma-Aldrich) in PBS for 10 min, followed by blocking solution composed of 10% normal goat serum (Abcam), 1% BSA (Research Products International), 0.3 M glycine (Sigma-Aldrich) and 0.1% Tween 20 (Sigma-Aldrich) in 10 mL DPBS (Corning) for 1 h at room temperature. ZO-1 monoclonal antibody (Invitrogen), Alpha Tubulin (TUBA4A) mouse monoclonal antibody (Origene) and MUC5AC monoclonal antibody (Invitrogen) were each diluted to 5 mg/mL in blocking solution. The samples were incubated with ZO-1, TUBB4A and MUC5AC overnight at 4 °C and then incubated with Alexa 488-F(ab’)2 fragment of goat anti-mouse IgG (Invitrogen) for 2 h at room temperature. At the same time, nuclei were stained with DAPI solution (1 mg/mL) for 2 h at room temperature. The cells were thoroughly washed three times for a total of 3 min between each procedure. Images were captured using a confocal microscope (Zeiss LSM880).

ALI samples were then characterized with four major epithelial cell markers, namely basal, goblet, club, and ciliated cells. They were embedded in OCT (Sakura Finetek) and snap frozen in liquid nitrogen. The frozen samples were cut at 8 µm thickness, air dried on Superfrost plus slides, then fixed with 4% PFA (15 min) and permeabilized with 1X PBS/0.1% Triton-X-100 (15 min). Tissue sections were treated with Fc Receptor Blocker, followed by Background Buster (Innovex Bioscience). The sections were stained with appropriate primary antibodies for 1 h followed by secondary antibodies for 30 min at room temperature in 1X PBS/5% BSA (bovine serum albumin)/0.05% saponin (**Table S1**). For panels including pure and conjugated antibodies, an additional step of blocking with 1/20 normal mouse serum in 1X PBS (15 min) was used (**Table S1**). Antibody validations were performed using isotype controls. Finally, sections were counterstained with 1 mg/mL of 4’,6-diamidino-2-phenylindole (DAPI) and 0.1 nmol units/ml of Phalloidin ATTO647N and mounted with Fluoromount-G (Thermo Fisher Scientific). The slides were acquired using a confocal (Leica SP8) (for high resolution images) or widefield microscope (Leica Thunder) (for histocytometry quantification based on immunofluorescent scans). Microscope acquisition was performed using LAS X software (Leica) and then image analysis was conducted using Imaris software (Oxford Instruments) (Bitplane).

### 2.7 ZO-1 quantitative analysis

With the obtained ZO-1 images, ZO-1-to-ZO-1 distance was quantitatively evaluated to represent how dense tight junctions were formed for different groups at Days 14 and 28. Using ImageJ Fiji software (National Institutes of Health), a single line was drawn on the fluorescent images to acquire the intensity profile. Then, peaks of the intensity profile were identified. Subsequently, the position of each peak was converted to the actual scale in μm and the distance between two peaks was measured.

### 2.8 Quantification of hNEC differentiation markers

After acquiring cilia and MUC5AC images, the area of fluorescence was quantified using ImageJ Fiji. For the cilia area, a region of 10^5^ mm^2^ was selected and analyzed to represent the area in absolute value, and for the MUC5AC area, the area was represented as a percentage.

### 2.9 Transepithelial electrical resistance (TEER) measurements

TEER values were measured with a STX2/chopstick electrode across the epithelial layer in Transwell inserts using an EVOM2 epithelial voltage meter (World Precision Instruments). According to the manufacturer’s instructions, electrodes were sterilized and adjusted before the measurement. To eliminate excessive mucus, cells were manually washed once with pre-warmed DPBS prior to TEER measurements. Subsequently, pre-equilibrated ALI medium (Stemcell Technologies) was added to the apical (0.25 mL) and basolateral (1 mL) chambers. Before reading TEER values, the monolayers were given time to reach a stable potential for ∼5 min inside the incubator. The sample resistance was calculated by subtracting the values from the cell-free Transwell insert’s resistance. Resistance of each Transwell insert in units of Ω was calculated using a previously described technique [36].

### 2.10 Calculation of ciliary beating frequency (CBF)

After 4 weeks of ALI culture, ciliary beating of hNECs was examined using a recently described technique [37]. The inserts were washed three times with prewarmed DPBS to eliminate the mucus. CBF was recorded as a series of images using the Axio Zoom microscope. The images were taken at a sample interval of 1 ms, a frame rate of 100 frames/s and 40x microscope magnification. A series of at least 512 or 1024 images was captured over a period of 10 s. For subsequent retrieval and analysis, each set of frame-by-frame photos was saved in a TIF type file. Three randomly chosen areas of a single insert were captured. Over the recording time, the pixel intensities of a particular area of interest (ROI) were collected, and the data were used for fast Fourier transformation (FFT). Using Equation 1, CBF was determined by counting a certain number of distinct cilia beat cycles:

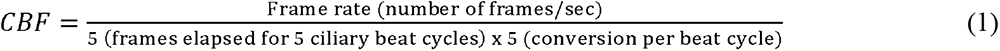

### 2.11 Permeability study

The permeability study was performed for all groups using a 4-kDa fluorescein isothiocyanate (FITC) Dextran marker (Sigma-Aldrich, CAT:46944-100MG-F). Dextran permeability assay was performed for samples at Day 28. Dextran (10 mg/ml) was applied to the apical part of Transwell inserts for cell-laden and blank samples and then all inserts were incubated in the incubator. Media was collected from the basal compartment every 60 min and loaded into 96-well plates (Greiner Bio) for 3 h. Using a microtiter plate reader (BioTek Instruments Inc.), Dextran content in the base samples were calculated using a 492-nm-excitation wavelength.

### 2.12 scRNA-seq dissociation

Inserts were washed with PBS, both apically and basally. They were then incubated twice with 0.05% trypsin/EDTA, apically and basally. 2% FBS and 2mM EDTA in PBS was used to inactivate trypsin. Following each incubation with trypsin, cells were collected by gently pipetting up and down without scratching the membrane with the pipette tip. Cells were washed with PBS and then incubated in a Dispase I/DNase I solution. The samples were then filtered through a pre-wetted 30 mm cell strainer. Single cells were washed and suspended in PBS containing 0.04% BSA and immediately processed as follows. Cell viability was assessed on a Countess II automated cell counter (ThermoFisher) and up to 10,000 cells were loaded onto a single lane of a 10X Chromium X. Single cell capture, barcoding and library preparation were performed using the 10X Chromium platform version 3.1 NEXTGEM chemistry and according to the manufacturer’s protocol (#CG000388) [38]. cDNA and libraries were checked for quality on Agilent 4200 Tapestation and ThermoFisher Qubit Fluorometer, quantified by KAPA qPCR, and sequenced at 16.6% of an Illumina NovaSeq 6000 S4 v1.5 200 cycle flow cell lane, with a 28-10-10-90 asymmetric read configuration, targeting 5,000 barcoded cells with an average sequencing depth of 80,000 read pairs per cell. Illumina base call files for all libraries were converted to FASTQs using bcl2fastq v2.20.0.422 (Illumina) and FASTQ files associated with the gene expression libraries were aligned to the GRCh38 reference assembly with v32 annotations from GENCODE (10x Genomics GRCh38 reference 2020-A) using the version 6.1.2 Cell Ranger multi pipeline (10x Genomics).

### 2.13 scRNA-seq analysis

Seurat was implemented for analysis of scRNA-seq data in this study. Seurat (version 4.2.1) was performed in R and was applied to all datasets. For downstream analyses, we kept cells which met the following filtering criteria per biological replicate per condition: >100 and <5000 genes/cell, >80% number of genes per unique molecular identifier (UMI) for each cell and <15% mitochondrial gene expression.

To select highly variable genes (HVGs) for initial clustering of cells, we performed FindVariableFeatures function on the normalized data for all genes included in the previous step. For clustering, we used the function FindClusters that implements Shared Nearest Neighbor modularity optimization-based clustering algorithm on 20 Principal components (PC) with a resolution of 0.8. Nonlinear dimensionality reduction methods, namely t-distributed stochastic neighbor embedding (tSNE) and uniform manifold approximation and projection (UMAP), were applied to the scaled matrix for visualization of cells in two-dimensional space using 20 PC components. The marker genes for every cluster compared with all remaining cells were identified using the FindConservedMarkers function.

R package Monocle3 (version 1.3.1) was used to reconstruct the epidermal cell (basal, suprabasal, club and goblet cells) developmental trajectory. To begin with, monocle3 converts the Seurat object into a monocle3 object using the as.cell_data_set function of the SeuratWrappers. Next, because partitions were high level separations of the data, we performed trajectory analysis on each partition separately. In addition, monocle3 added the trajectory graph to the cell plot via learn_graph() function, and the basal cell was defined as the root state argument based on the annotated gene for trajectory analysis. Finally, we exported this data to the Seurat object and visualized.

### 2.14 Viral Infection

Influenza virus (PR8-GFP, with GFP fused to NS1; A/PR8/34 (H1N1) [39]) was obtained from Dr. Adolfo García-Sastre (Icahn School of Medicine at Mount Sinai). Infectious stocks were prepared on Madin-Darby canine kidney (MDCK) cells and supernatants were centrifuged and filtered (0.45 mm). Viral titers were determined by plaque assay on MDCK cells. Aliquots were stored in a secured -80 °C freezer until further use.

Prior to infection, the apical surface of the ALI cultures was washed 1-3 times with PBS (1X) for 15 min at 37°C to remove the mucous. Then, the ALI samples were apically exposed to 25 mL of virus solution. The virus was diluted in Pneumacult ALI media serum-free for a final concentration of 2.5 × 10^5^ pfu PR8-GFP. Following 24 h incubation, 100 mL of PBS was added to the apical side of the ALI cultures for 15 min at 37 °C and harvested in an Eppendorf tube. Flow cytometric monitoring of virus infection was analyzed using recently described studies [40,41].

### 2.15 Histocytometry

To proceed to histocytometry, we adopted the Germain methodology [42]. Briefly, three consecutive sections, 8-mm thick, were acquired and the intensity mean for cell populations positive for DAPI (Y axis) and viral GFP (X axis) were generated in Imaris 9.7 (Oxford Instruments) using the Channel Arithmetics Xtension prior to running surface creation to identify DAPI-GFP cells in images. Statistics were exported for each surface and imported into FlowJo v10.3 (BD Bioscience) for image analysis.

### 2.16 Statistical analysis

The data were displayed by using GraphPad Prism 8 (GraphPad Software) and statistical analysis was carried out using SPSS 28 software (SPSS, Inc.). One-way ANOVA with post-hoc Tukey tests was used for comparison of the differences among multiple groups, which were indicated as *\*p* < 0.05, *\*\*p* < 0.01, and *\*\*\*p* < 0.001.

## 3. Results

### 3.1 DBB of the Nasal Epithelial Tissue

In our experiments, we first verified the biocompatibility of DBB using LIVE/DEAD staining, which captured live (green) and dead (red) cells at Day -8 and -2 (**Figure 2A**). A day after bioprinting (Day -8), all groups showed high cell viability (over 90%), and such a viability level was maintained for a week (Day -2) after bioprinting (**Figure 2B**). These results indicate that DBB was compatible with hNECs as cells were able to attach, grow, and differentiate afterwards.

**Figure 2.**
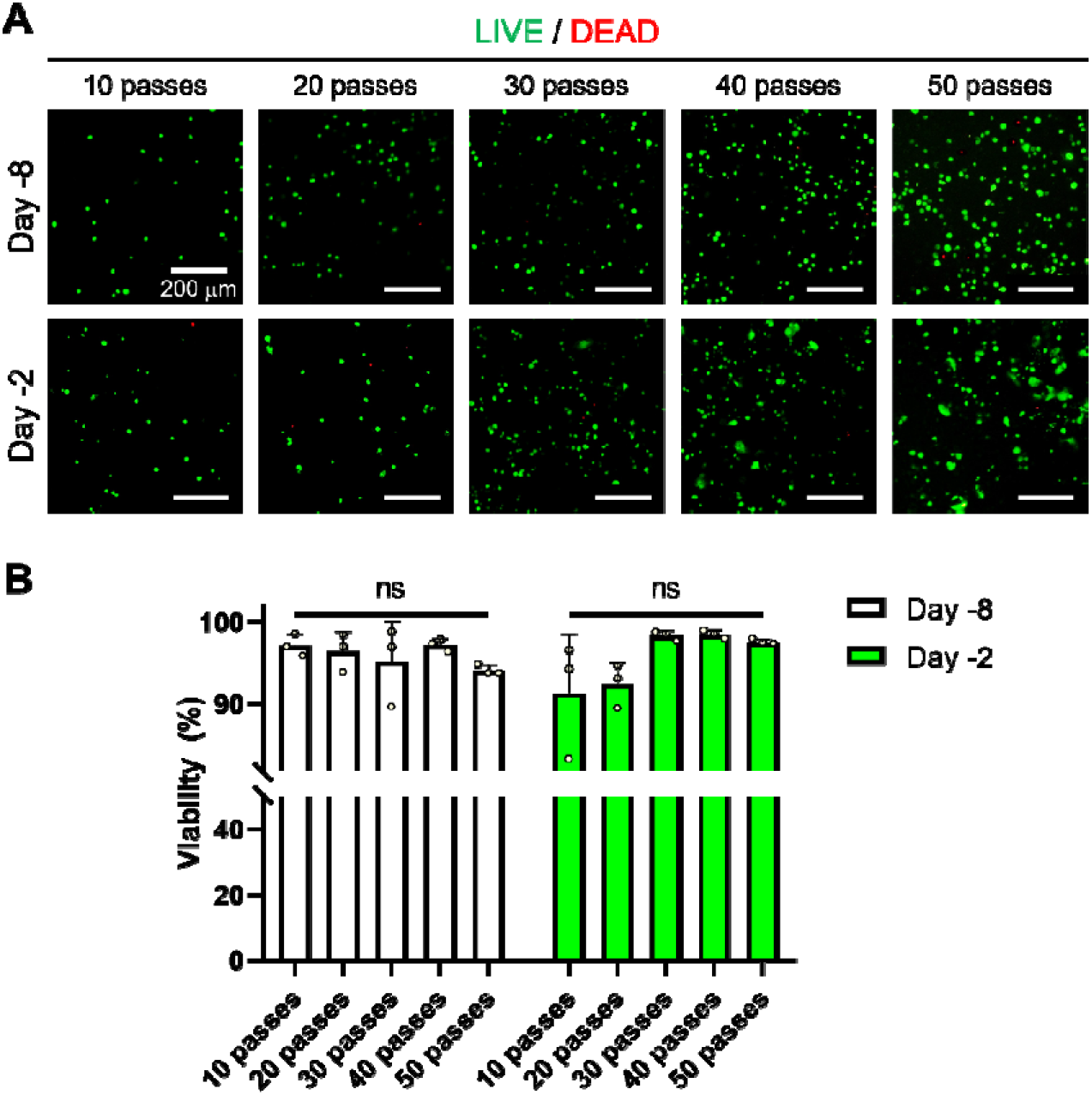
Cell viability measurements using the LIVE/DEAD assay. **(A)** Representative fluorescent images of hNECs at different densities at Day -8 and Day -2, and **(B)** cell viability (%) for all bioprinted groups including 10-, 20-, 30-, 40- and 50-pass groups (*n*=3; ns denotes ‘not significant’).

After confirming the cell viability, cell growth was tracked and captured for a more comprehensive analysis. **Figure S1** demonstrates optical images of hNECs at Day -9 (a day after DBB), Day 0 (the end of liquid-liquid culture), and Day 28 (the end of the differentiation process). At Day -9, hNECs were identified in all groups including manual seeding and bioprinted (with different number of pass groups). However, by the end of the liquid-liquid culture (Day 0), it was clearly shown that the cell population did not further expand for 10- and 20-pass groups. To maintain cellular activities, hNECs may require cell-cell interactions, but 10 and 20 passes did not provide the required cell density for these cell-cell interactions. Hence, while other samples started to form cell layers from Day 0, 10- and 20-pass groups revealed single cell configuration until the end of differentiation process (Day 28). For the rest of the study, 10- and 20-pass groups were thus not considered, and experiments were carried out using the samples of manual seeding (control) and bioprinted (with the number of passes over 20) groups.

To confirm the tight junctions, ZO-1, a classical scaffold protein maintaining cell-cell adhesion found in stable tissues, was imaged (**Figure 3A**), and ZO-1-to-ZO-1 distance was measured at two different time points, Day 14 (middle of the differentiation process) and Day 28 (end of the differentiation process) [43]. We observed a significant difference between the samples from manual seeding and bioprinted groups at Day 14 as shown in **Figure 3B**. At Day 28, when the differentiation process was completed, 30-pass and manual seeding groups became similar but there was a significant difference between manual seeding and 40- and 50-pass groups (**Figure 3C**). Overall, we showed that tight junctions can be modulated by the number of passes and distributed more uniformly when DBB was applied.

**Figure 3.**
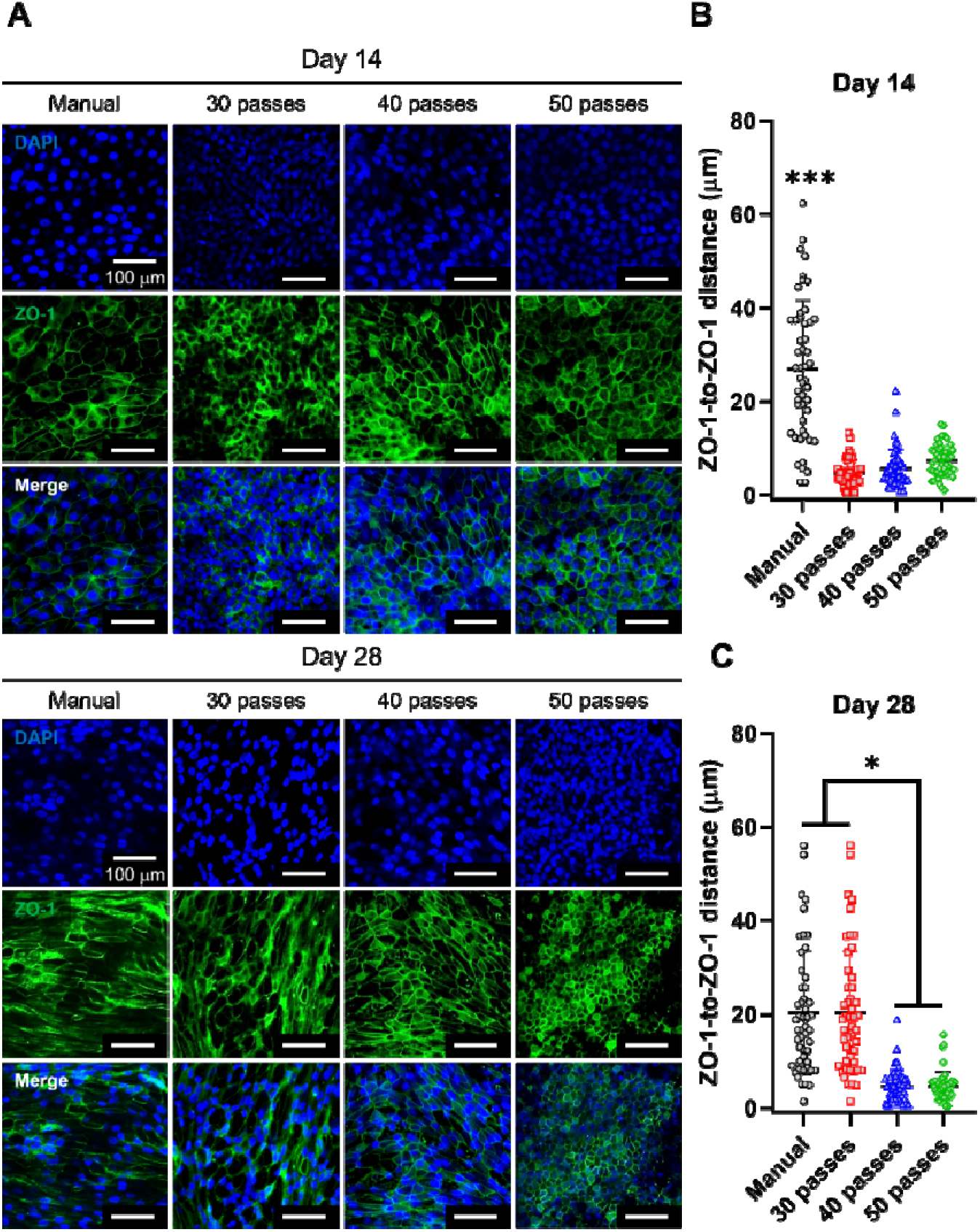
Evaluation of tight junctions in nasal epithelium using ZO-1 staining. **(A)** Confocal images of ZO-1 staining of hNECs in ALI culture of manual (control) and bioprinted (30-, 40- and 50-pass) groups at Weeks 2 and 4. Characterization of ZO-1 staining with respect to ZO-1-to-ZO-1 distance at Week **(B)** 2 and **(C)** 4 (*n*=3; p*<0.05 and p***<0.001).

Since the differentiation process was completed at Day 28, cilia and MUC5AC positive area were evaluated (**Figure 4**). According to our calculations, the ratio of cilia per 10^5^ mm^2^ in the manual seeding group was much less compared to that in bioprinted groups (**Figure 4B**). This leads us to conclude that the bioprinted groups had a higher density of ciliated cells implying accelerated cell development and differentiation. On the other hand, there was a significant difference between 30- and 50-pass groups in terms of MUC5AC staining (**Figure 4D**). These results indicated that the 50-pass group was more differentiated in terms of the presence of goblet cells.

**Figure 4.**
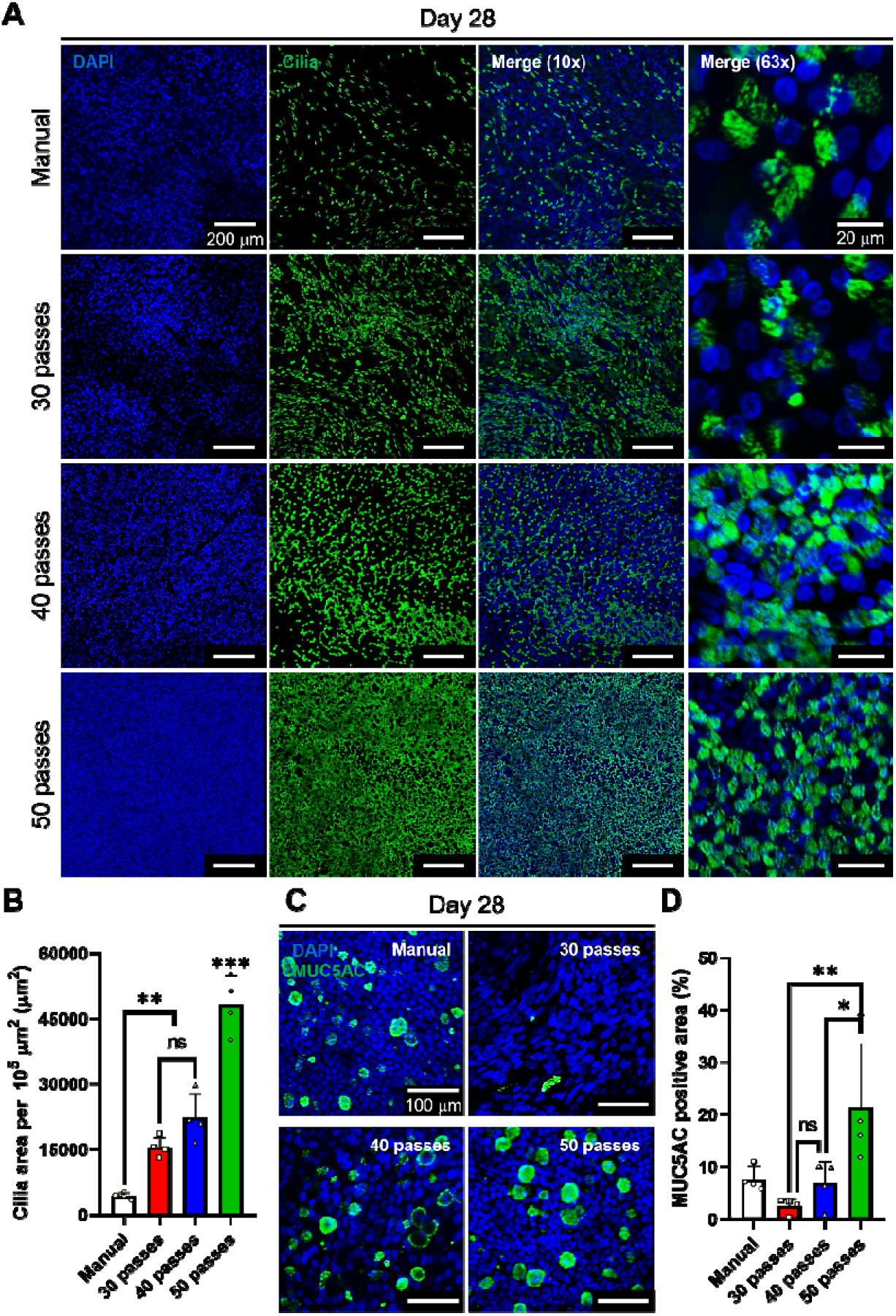
Evaluation of cilia formation and mucus secretion of nasal epithelium fabricated via manual seeding or bioprinting using a-Tubulin (cilia) and MUC5AC staining. **(A)** Confocal images of cilia staining of hNECs at Day 28 of ALI culture. **(B)** Characterization of cilia expression using cilia area per 10^5^ μm^2^ at Day 28 of ALI culture. **(C)** Confocal images showing MUC5AC staining of hNECs at Day 28 of ALI culture and **(D)** related characterization of mucus production using MUC5AC positive areas (%) (*n*=3; p*<0.05, p**<0.01 and p***<0.001; ns denotes ‘not significant’).

### 3.2 Functional analysis of the bioprinted nasal epithelium

In this study, to verify the barrier function of the bioprinted nasal epithelium, TEER measurements were performed for the first 4 weeks of ALI culture (**Figure 5A**). The mean TEER value after a week in ALI was 1003 Ω × cm^2^ for the 50-pass group, which was higher than the average TEER of 971 Ω × cm^2^ for the manual group (p = 0.0003). After 28 days of differentiation under ALI, the TEER value was 252 Ω × cm^2^ for the manual group and 255 Ω × cm^2^ for the 50-pass group. Overall, all groups exhibited a similar TEER trend over time and the TEER measurements at Day 28 were similar. In addition, the tightness of the epithelium was confirmed by a permeability study (**Figure 5B**). The flow of 4 kDa FITC-labeled Dextran from the upper chamber to the lower chamber was significantly reduced for the 28-day ALI cultured samples compared to the cell-free synthetic Transwell membranes (control). The corresponding reduction in flux rate was similar among manual, 30-, 40- and 50-pass groups, but decreased as the number of passes increased. The Dextran influx results supported the findings with TEER measurements that our samples allowed the formation of a tight epithelium under the same differentiation conditions.

**Figure 5.**
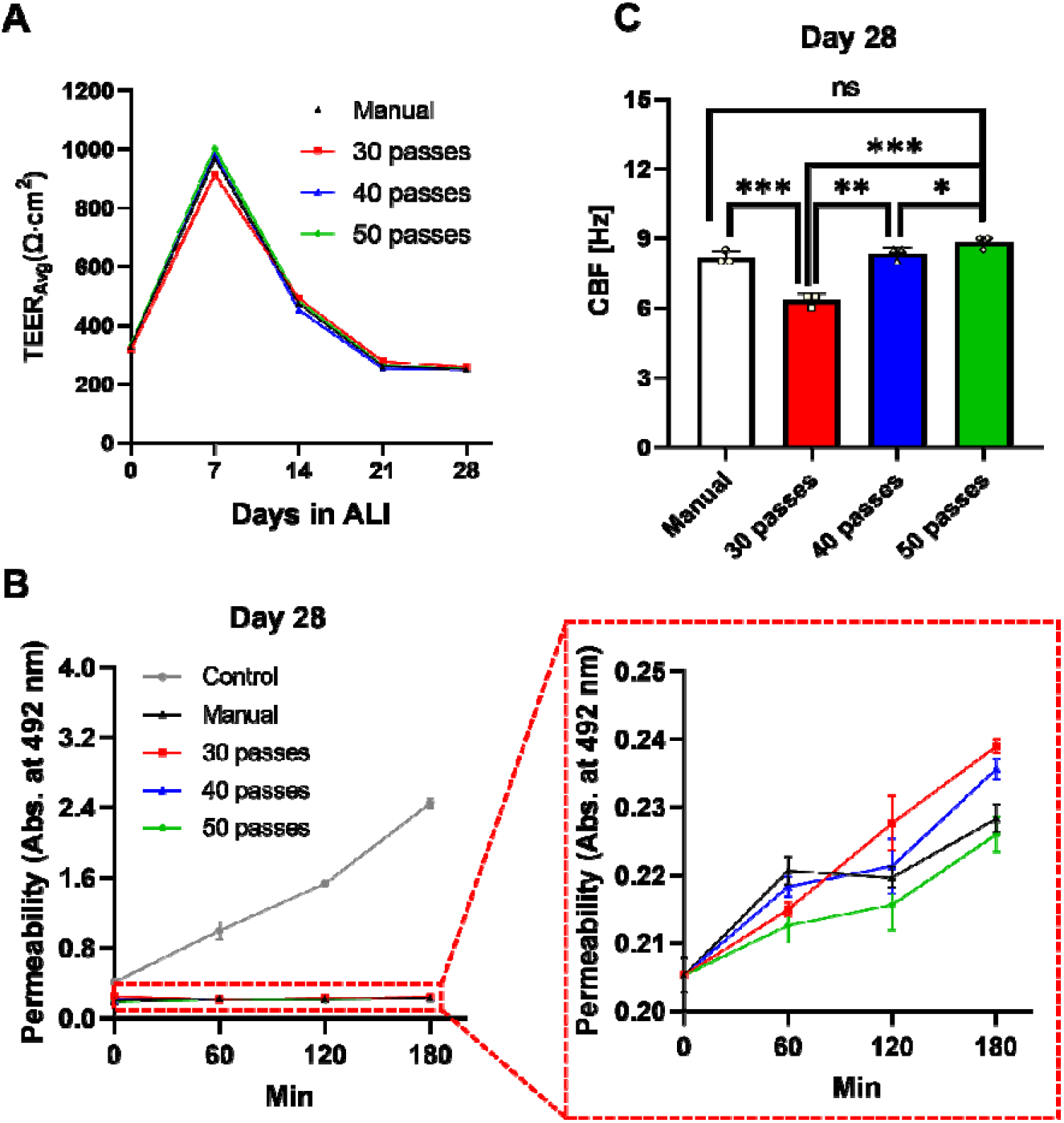
Functional analysis of nasal epithelial tissues, including manual and bioprinted (30-, 40- and 50-pass) groups. **(A)** Measurement of TEER values during the 28-day differentiation. **(B)** Comparison of Dextran permeability and **(C)** CBF at Day 28 (*n*=3; p*<0.05, p**<0.01 and p***<0.001; ns denotes ‘not significant’).

A functional nasal epithelium in-vivo is characterized by synchronized beating of cilia. To confirm the ciliary function of ALI cultured hNECs at Day 28, CBF was determined by microscopic recording combined with image analysis (**Figure 5C**). The median CBF of the 50-pass group was 8.83 Hz, which was in a comparable range with respect to the manual group (8.17 Hz). The median CBF for the 40-pass group was 8.34 Hz while the median CBF for the 30-pass group was 6.34 Hz. Overall, the CBF values increased as the number of passes increased for the bioprinted groups (**Video S2**).

As the functional analysis revealed similar TEER and permeability measurements for all bioprinted groups, we proceed with the 30-pass group for the rest of the study.

### 3.3 Histological and scRNA-seq analysis

Next, for the 30-pass group, we performed a histological study using immunofluorescence imaging. We used cell-type specific markers to visualize and quantify the presence of major epithelial cell types (**Figure 6**). Bioprinted nasal epithelium effectively recapitulated the in-vivo upper airway pseudostratified ciliated columnar epithelial architecture with the presence of ciliated, club, goblet and basal cells [44,45] . These data indicated that hNECs were well-differentiated with mainly consisting of basal cells. The presence of functional mucus-producing cells was further supported by the staining of mucus-containing glycoproteins with MUC5AC. A high amount of mucus staining was observed in 28-day ALI culture as the differentiation was attained at the highest level at Day 28.

**Figure 6.**
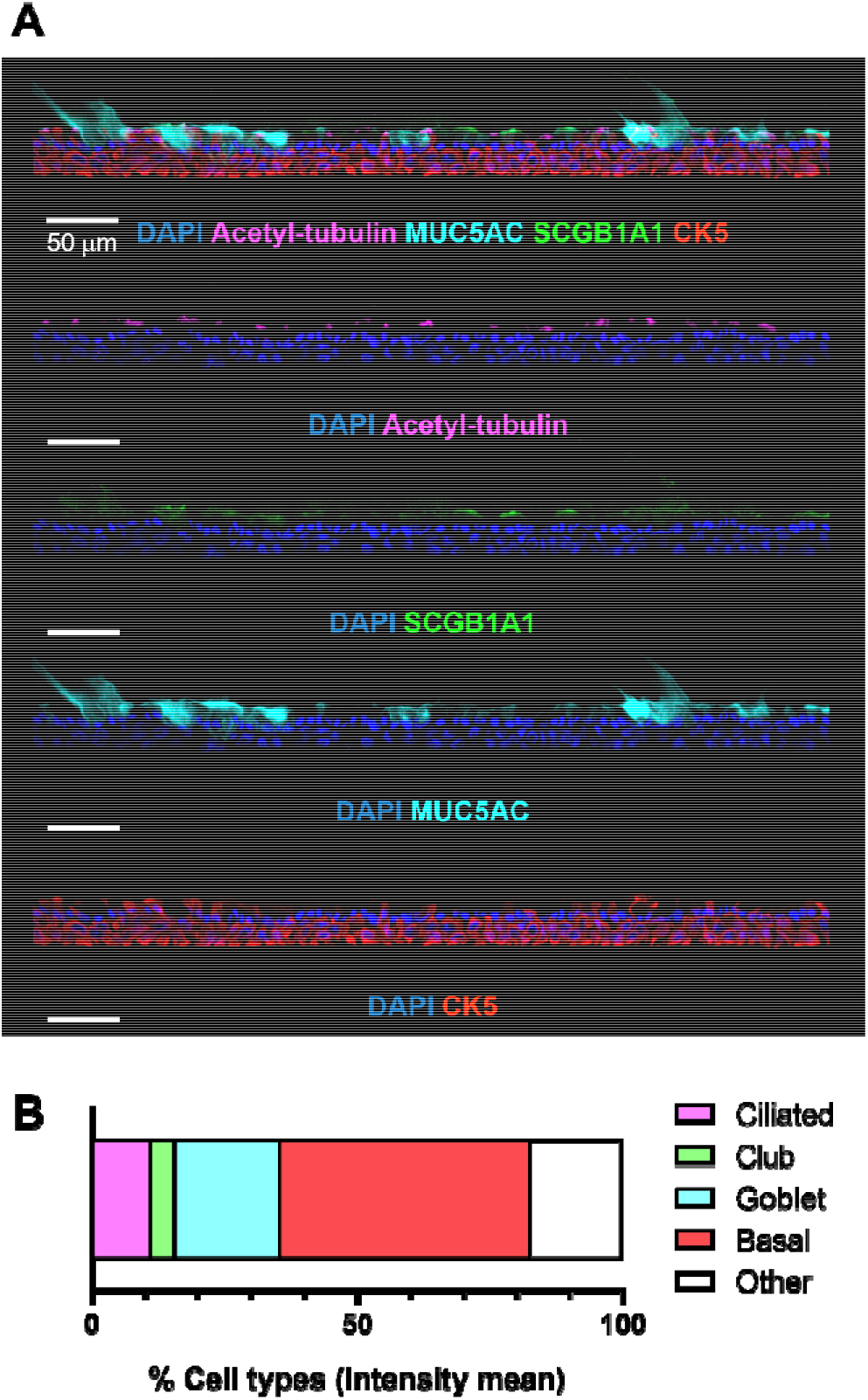
Cellular composition of the bioprinted (30-pass) nasal epithelium. **(A)** Representative immunofluorescent images of differentiated hNECs characterized by major epithelial cell markers: basal (CK5, red), goblet (MUC5AC, cyan), club (SCGB1A1, green) and ciliated cells (acetylated α-tubulin, magenta) and nuclei (DAPI, blue). **(B)** Quantification of epithelial cell types in nasal ALI cultures by histocytometry (*n*=1).

Next, we performead scRNA-seq analysis for the 30-pass group. We utilized a total of 7315 cells from the bioprinted nasal cultures with single-cell RNA sequencing (scRNA-seq). After performing quality control using Seurat, we assembled RNA sequencing data from 3,288□cells for clustering analysis (**Figure S2A**). This demonstrated 5 nasal tissue cell populations that were visualized by uniform manifold approximation and projection (UMAP) embeddings (**Figures 7A and 7B**). Clusters were identified by specific gene expression (**Figure S2B**, **Table S2**), determining basal, suprabasal, club, goblet, and multiciliated cells (**Figure 7A**). Meanwhile, we found basal cells underwent a bidirectional differentiation into goblet cells and multiciliated cells (**Figure 7C**), which is identical to known human nasal cell development [46]. During differentiation, basal cells gradually lost expression of basal markers (*DLK2, KRT14, LAML3*) and simultaneously started to express suprabasal markers (*FABP5, KRT13*) (**Figure 7D**). This biological observation allowed us to generate a pseudo time trajectory inferred by Monocle3 (see Methods) and reconstructed a putative basal-superbasal-club-goblet cell differentiation trajectory (**Figure 7D**). As expected, Monocle3 inferred the differentiation to start with a group of highly expressed *KRT5* basal cells and tracked the differentiation with decreasing expression levels of *KRT5* (**Figure 7D**). In short, with the scRNA-seq, we identified five cell populations in our nasal samples, and the trajectory analysis hinted that the basal cells had the potential to differentiate into goblet cells and multiciliated cells.

**Figure 7.**
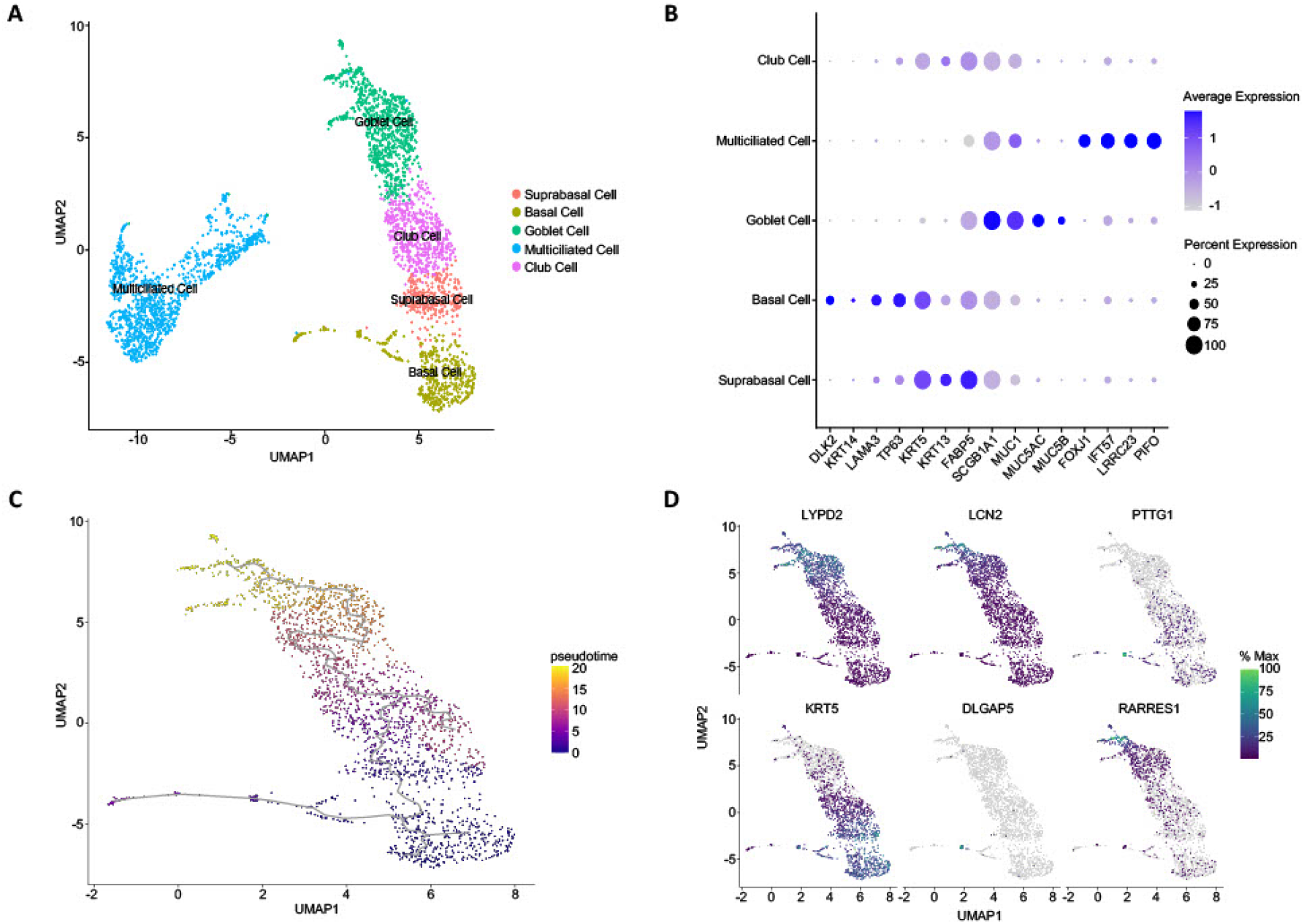
Defining bioprinted nasal cells populations and differentiation trajectory using scRNA-seq. **A)** UMAP plot visualization of 3288 single cells isolated from bioprinted nasal epithelium sample SC2200969 that passed quality control metrics using Seurat. Putative cellular communities are presented on the right. **(B)** Dotplot heatmap of relative gene expression of known nasal epidermal marker genes split by cell clusters using Seurat. **(C)** UMAP plot showing pseudotime inference of bioprinted nasal epithelium sample SC2200969. **(D)** UMAP plot overlays showing selected gene expression distribution across clusters for pseudotime inference.

### 3.4 Viral infection of the bioprinted nasal epithelium

Lastly, we determined if these high-throughput bioprinted nasal epithelial cultures were permissive to respiratory viral infection. For that, we exposed ALI cultures to influenza virus strain A/PR8/34 carrying GFP (2.5 × 10^5^ pfu) or serum-free Pneumacult ALI media, where the uninfected mock group was used as a control. Following a 24-h incubation, we harvested the ALI cultures and subsequently performed immunofluorescence staining (**Figure 8A**). We detected the presence of virus through GFP and visualized the structural integrity of the cultures using Phalloidin (F-Actin) staining. The presence of GFP signal indicated active infection as GFP was a gene reporter for non-structural protein 1 (NS1). The data demonstrated the functionality of our model as shown in **Figure 8B**, where ∼11% of the cells were infected.

**Figure 8.**
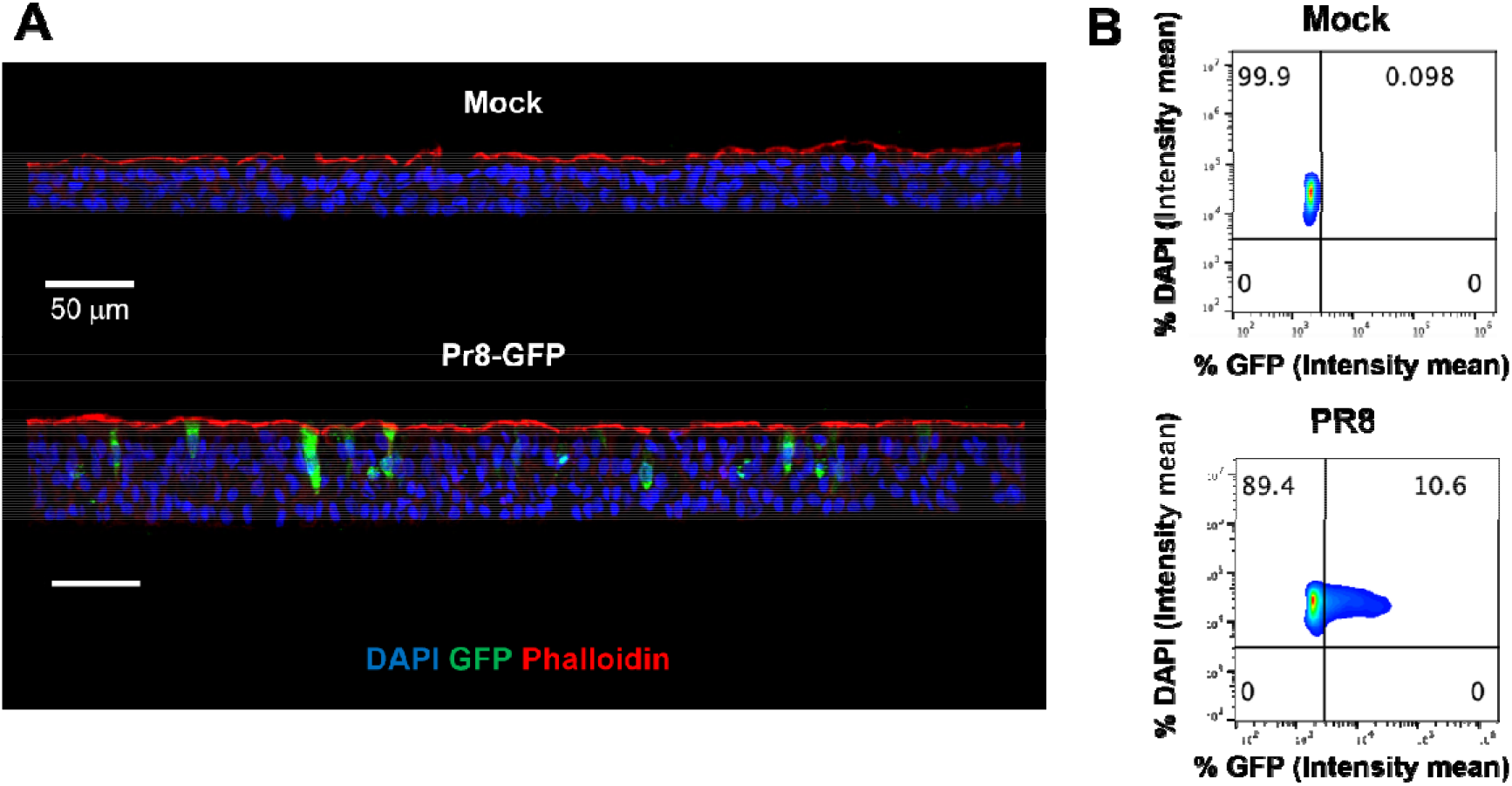
Bioprinted nasal epithelial tissue samples permissive to GFP^+^ influenza infection. **(A)** Representative immunofluorescent images following 24 h exposure to uninfected control (mock) or influenza virus (2.5 × 10^5^ pfu GFP^+^ PR8); nuclei (DAPI, blue), Phalloidin (actin filament, red), and GFP^+^ influenza virus to reveal effective viral replication (GFP, green). **(B)** Representative histocytometry dot plots of mock and GFP^+^ PR8 conditions, showing the intensity mean for cell populations positive for DAPI (Y axis) and virus reporter GFP (X axis).

## 4. Discussion

To better understand the role of respiratory epithelial cells in protecting the body and regulating the immune response, it is important to study these cells in a way that mimics the in vivo situation. This can involve looking at differences among epithelial cells from diseased populations or studying the effects of external factors on these cells [47,48]. Human airways contain many distinct types of cells, including ciliated, goblet, and basal cells, and studying these cells in an in-vivo like setting can provide valuable insights [49]. Using an in vitro model of the human nasal tissue, researchers can study specific differences in nasal epithelial cells related to different diseases, investigate the underlying mechanisms of these diseases, and control exposure to air pollutants [50–52]. Additionally, researchers can examine cell-cell interactions in a controlled environment by growing these cells on tissue culture inserts [53,54].

In this study, we developed a method for bioprinting and investigating the human nasal epithelial tissue, which has been rarely studied compared to the bronchial epithelial cultures. There are multiple advantages of studying nasal epithelial tissue instead of its bronchial counterpart [55–59] . First, nasal epithelial cells are often the first to be exposed to environmental stressors, such as pollutants or allergens, making them valuable models for studying the effects of these stressors on airway epithelial cells. Secondly, it is not always possible or appropriate to perform bronchoscopy to obtain lower airway cells in individuals with certain pre-existing diseases. Instead, the less invasive method of brushing the nasal turbinate can be used. Additionally, nasal biopsies can be done multiple times without significant side effects, making them useful for repeated studies [60–63] .

There are various commercially available cell culture systems of pre-differentiated human epithelial cells, such as the EpiAirway model from MatTek, Epithelix Sárl from Switzerland, and Clonetics from Lonza, which are known to be stable (reproducible using standardized protocols and operating procedures and meeting quality standards) [64–67]. However, these systems can be expensive, and the number of samples is limited. Additionally, it can be difficult to control the characteristics of the donors, such as age, gender, and disease status, when using commercially available cultures. On the other hand, using freshly obtained nasal epithelial cells from superficial brush or scrape biopsies allows the researcher to control the characteristics of the donors and to recall the same volunteers for follow-up studies. Bioprinted epithelial tissue culture models are a promising alternative to the epithelial tissue models produced using manual approaches, such as aforementioned tissue models on the market. Bioprinting has several advantages compared to the manual approaches: it can be high-throughput and precise. In addition, it is a highly automated process, enabling the rapid production of large numbers of tissue constructs. This allows for high-throughput experimentation, such as large-scale drug and microbial screening [68,69]. Furthermore, it enables precise control over the spatial arrangement of cells, which is significant for creating functional tissue structures. This is particularly useful for creating tissue models that mimic the structure and organization of native tissues [70,71]. Besides, bioprinting allows for the homogeneous distribution of cells within a tissue construct ensuring that the cells are evenly distributed and that the tissue has a homogeneous cellular composition and tight junctions, which might be essential to create tissues with barrier function [72,73]. Lastly, bioprinting can help with the reduction in the costs associated with manual cell culture techniques (less media, less equipment, etc.) as it allows for the rapid and efficient production of primary tissue cultures [74].

The optimization of the bioprinting process was essential to ensure that the bioprinted nasal epithelium was functional and mimics the structure of its native counterpart. Although there are multiple studies on hNECs and optimization of their ALI culture [13] for viral infection [75] or drug screening [76,77], this is the first study to use bioprinting of hNECs. With this study, we have shown that bioprinting much lower number of cells with respect to the traditional manual seeding approach resulted in nasal epithelium with a higher degree of differentiation. Our study confirms that despite the differences in the initial cell numbers, the proportion of ciliated and goblet cells were comparable to previously reported studies. The proportion of ciliated cells in culture conditions was consistent with the proportion reported in normal, healthy human airway epithelium (50-70%) reported previously [78]. However, the proportion of goblet cells found in our study was higher than the typical percentage found in adult human airway epithelium, which was reported to be up to 25% of cells [79]. The authors noted that the proportion of goblet cells found in that study was also higher compared to the previous studies of the authors on cultures derived from newborn and 1-year-old infants [80]. Even though the reason for this discrepancy is unclear, it may stem from donor- or age-specific factors.

Permeability data, particularly for multi-layered structures, is essential in determining the applicability of permeability values obtained from in-vitro experiments to clinical studies. It is important to note that various cell layers within human epithelial tissues possess distinct permeability properties, and the thickness of the culture model or tissue should be considered and controlled when comparing TEER or permeability values between different in-vitro models [81]. Nasal epithelial tissue, maintained for 28 days, exhibited high levels of cell shedding at the ALI interface, which may have been responsible for the increased Dextran permeation. This is supported by the TEER measurements, which showed that TEER of the nasal epithelium decreased as the cells were cultured for longer periods of time. Together, these findings suggest that cell shedding at the ALI interface is likely a major contributor to the observed increase in Dextran permeation. TEER values are a measure of the integrity of the epithelial cell monolayer, which is a crucial aspect of tissue culture [36,82]. TEER values in the presented study were higher than those reported in previous studies of human-adult [83] and pediatric nasal epithelial cultures [84]. The maximum TEER values reported in those studies were ∼400 ohm×cm^2^ after 9 days and ∼150 ohm×cm^2^ after 14 days. The values indicate that the epithelium in the current study had a higher integrity, meaning that the cells were more tightly packed and had better barrier function, than the cells reported in these earlier studies. Other than TEER, CBF plays a critical role in the mucociliary clearance process; however, the mechanisms responsible for regulating CBF are yet to be fully understood [85]. In our study, the mean CBF for all samples ranged from 6.34 to 8.83 Hz, slower than the typical CBF of healthy cilia in the human respiratory mucosa. According to the literature, the CBF of healthy cilia in the human respiratory mucosa is typically around 11 to 16 Hz [86,87]. This suggests that the cilia in our samples may not be functioning as perfect as healthy cilia. The deviation of the mean CBF from the literature values is quantified by the standard deviation of 5.4 Hz, which indicates that some of the samples had CBF within the range of healthy cilia and some of them were lower. On the other hand, the results of our study pertaining to CBF values are in agreement with recent studies on human nasal epithelial cultures exposed to repetitive preservatives, suggesting consistency and potential applicability of the current study’s findings to understand the effects of preservatives on CBF [88,89].

One of the advantages of the presented approach is that it allows for the creation of the nasal tissue that closely mimics the structure and function of the native nasal epithelium, which can be useful for studying the behavior of nasal epithelial cells in response to different drugs or pathogens, as well as for developing new treatments for nasal conditions such as chronic sinusitis. Indeed, the still ongoing COVID-19 pandemic has shown the emergence of in vitro models, such as ALI cultures, to study the response to SARS-CoV-2 virus [90,91]. Furthermore, the evolution of the virus with the emergent variants of concern (VOC), has shown that in vitro ALI cultures from upper airway epithelium, nasal and bronchial, are highly permissive to SARS-CoV-2 variants [44,45,92,93]. Personalized medicine is another potential application of the bioprinted nasal epithelium. By bioprinting patient-specific hNECs, it may be possible to create personalized treatments that are more effective and have fewer side effects. Despite the potential benefits, there are still challenges, such as biochemical properties, bioactivity and mechanical properties that need to be addressed before using such a system in clinical practice [94].

## 5. Conclusion

This study presents the development of a nasal epithelial tissue model using primary hNECs in a high-throughput manner via DBB. Progenitor hNECs were bioprinted with different numbers of passes ranging from 10 to 50, which were benchmarked against the commonly used manual seeding approach. After bioprinting, hNECs showed high cell viability regardless of the pass number and we demonstrated that tight junctions, mucus secretion, and ciliary beat frequency can be modulated by the number of passes. Bioprinted hNECs resulted in a higher degree of differentiation compared to manual seeding, even though the bioprinted cell density per insert was half or one third that of the manually seeded cell density per insert. Five cell populations were identified in the bioprinted tissue mimicking the native nasal epithelium and the trajectory analysis revealed that the bioprinted hNECs had the potential to differentiate into goblet and multiciliated cells. The presented method may inspire a shift in current hNEC practices and can be used for various applications such as infection studies, drug testing and disease modelling.

## Data availability statement

The single cell matrix is available from https://github.com/ohlab/SC2200969_Nasal30pass.

Raw fastq data available upon request.

## Acknowledgments

This research was primarily supported by the National Institute of Health Award #U19A142733. We gratefully acknowledge the contribution of the Single Cell Biology service, the Genome Technologies service, and cyberinfrastructure high performance computing resources at The Jackson Laboratory (JAX) for expert assistance with the work described herein. These shared services were supported in part by the JAX Cancer Center (P30 CA034196). JO was additionally supported by 1DP2GM126893-01, 1 R01 AR078634-01 and 5 R21 AR075174-02. MH was supported by T32HG010463. The Cure Cystic Fibrosis Columbus (C3) Epithelial Cell Core at Nationwide Children’s Hospital (NCH) provided primary human nasal epithelial cultures for this work, with the help of the NCH Biopathology Center Core and Data Collaboration Team. C3 is supported by a Cystic Fibrosis Foundation (CFF) Research Development Program grant (MCCOY17R2) and a CFF grant to the NCH Division of Pediatric Pulmonary Medicine (MCCOY19RO). The authors are also thankful to Dr. Adolfo García-Sastre and Michael Schotsaert from Icahn School of Medicine at Mount Sinai, New York, NY for providing, GFP^+^ PR8 virus.

## References

[1.] Geurkink N 1983 Nasal anatomy, physiology, and function J Allergy Clin Immunol 72

[2.] Harkema J R, Carey S A and Wagner J G 2006 The Nose Revisited: A Brief Review of the Comparative Structure, Function, and Toxicologic Pathology of the Nasal Epithelium Toxicol Pathol 34

[3.] Harkema J R 2015 Comparative Anatomy and Epithelial Cell Biology of the Nose Comparative Biology of the Normal Lung: Second Edition

[4.] Wanner A, Salathe M and O’Riordan T G 1996 Mucociliary clearance in the airways Am J Respir Crit Care Med 154

[5.] Antunes M B and Cohen N A 2007 Mucociliary clearance – A critical upper airway host defense mechanism and methods of assessment Curr Opin Allergy Clin Immunol 7

[6.] Kojima T, Go M, Takano K I, Kurose M, Ohkuni T, Koizumi J I, Kamekura R, Ogasawara N, Masaki T, Fuchimoto J, Obata K, Hirakawa S, Nomura K, Keira T, Miyata R, Fujii N, Tsutsumi H, Himi T and Sawada N 2013 Regulation of tight junctions in upper airway epithelium Biomed Res Int 2013

[7.] Siti Sarah C O, Shukri N M, Mohd Ashari N S and Wong K K 2020 Zonula occludens and nasal epithelial barrier integrity in allergic rhinitis PeerJ 8

[8.] Min H J, Kim T H, Yoon J H and Kim C H 2015 Hypoxia increases epithelial permeability in human nasal epithelia Yonsei >Med J 56

[9.] Huang Z Q, Liu J, Ong H H, Yuan T, Zhou X M, Wang J, Tan K sen, Chow V T, Yang Q T, Shi L, Ye J and Wang D Y 2020 Interleukin-13 Alters Tight Junction Proteins Expression Thereby Compromising Barrier Function and Dampens Rhinovirus Induced Immune Responses in Nasal Epithelium Front Cell Dev Biol 8

[10.] Riise G C, Andersson B, Ahlstedt S, Enander I, Söderberg M, Löwhagen O and Larsson S 1996 Bronchial brush biopsies for studies of epithelial inflammation in stable asthma and nonobstructive chronic bronchitis European Respiratory Journal 9

[11.] Muhlebach M S, Reed W and Noah T L 2004 Quantitative Cytokine Gene Expression in CF Airway Pediatr Pulmonol 37

[[12] Auerbach O, Stout A P, Hammond E C and Garfinkel L 1961 Changes in Bronchial Epithelium in Relation to Cigarette Smoking and in Relation to Lung Cancer New England Journal of Medicine 265

[[13] Müller L, Brighton L E, Carson J L, Fischer W A and Jaspers I 2013 Culturing of human nasal epithelial cells at the air liquid interface Journal of Visualized Experiments

[14.] Zhang H, Fu W and Xu Z 2015 Re-epithelialization: A key element in tracheal tissue engineering Regenerative Med 10

[15.] Derman I D, Singh Y P, Saini S, Nagamine M, Banerjee D and Ozbolat I T 2023 Bioengineering and Clinical Translation of Human Lung and its Components Adv Biol 2200267

[16.] Kabir A, Datta P, Oh J, Williams A, Ozbolat V, Unutmaz D and Ozbolat I T 2021 3D Bioprinting for fabrication of tissue models of COVID-19 infection Essays Biochem 65

[17.] Upadhyay S and Palmberg L 2018 Air-liquid interface: Relevant in vitro models for investigating air pollutant-induced pulmonary toxicity Toxicological Sciences 164

[[18] Lacroix G, Koch W, Ritter D, Gutleb A C, Larsen S T, Loret T, Zanetti F, Constant S, Chortarea S, Rothen-Rutishauser B, Hiemstra P S, Frejafon E, Hubert P, Gribaldo L, Kearns P, Aublant J M, Diabaté S, Weiss C, de Groot A and Kooter I 2018 Air-Liquid Interface in Vitro Models for Respiratory Toxicology Research: Consensus Workshop and Recommendations Appl In Vitro Toxicol 4

[[19] Zscheppang K, Berg J, Hedtrich S, Verheyen L, Wagner D E, Suttorp N, Hippenstiel S and Hocke A C 2018 Human Pulmonary 3D Models For Translational Research Biotechnol J 13

[20.] Papazian D, Würtzen P A and Hansen S W K 2016 Polarized Airway Epithelial Models for Immunological Co-Culture Studies Int Arch Allergy Immunol 170

[21.] Ozbolat I T 2016 3D Bioprinting: Fundamentals, Principles and Applications

[22.] Dey M and Ozbolat I T 2020 3D bioprinting of cells, tissues and organs Sci Rep 10

[23.] Ozbolat I T, Peng W and Ozbolat V 2016 Application areas of 3D bioprinting Drug Discov Today 21

[24.] Jin R, Cui Y, Chen H, Zhang Z, Weng T, Xia S, Yu M, Zhang W, Shao J, Yang M, Han C and Wang X 2021 Three-dimensional bioprinting of a full-thickness functional skin model using acellular dermal matrix and gelatin methacrylamide bioink Acta Biomater 131

[25.] Kim W J and Kim G H 2020 An intestinal model with a finger-like villus structure fabricated using a bioprinting process and collagen/SIS-based cell-laden bioink Theranostics 10

[26.] Madden L R, Nguyen T v., Garcia-Mojica S, Shah V, Le A v., Peier A, Visconti R, Parker E M, Presnell S C, Nguyen D G and Retting K N 2018 Bioprinted 3D Primary Human Intestinal Tissues Model Aspects of Native Physiology and ADME/Tox Functions iScience 2

[27.] Karamchand L, Makeiff D, Gao Y, Azyat K, Serpe M J and Kulka M 2023 Biomaterial inks and bioinks for fabricating 3D biomimetic lung tissue: A delicate balancing act between biocompatibility and mechanical printability Bioprinting 29 e00255

[28.] Horvath L, Umehara Y, Jud C, Blank F, Petri-Fink A and Rothen-Rutishauser B 2015 Engineering an in vitro air-blood barrier by 3D bioprinting Sci Rep 5

[[29] Berg J, Weber Z, Fechler-Bitteti M, Hocke A C, Hippenstiel S, Elomaa L, Weinhart M and Kurreck J 2021 Bioprinted multi-cell type lung model for the study of viral inhibitors Viruses 13

[30.] Shrestha J, Ryan S T, Mills O, Zhand S, Razavi Bazaz S, Hansbro P M, Ghadiri M and Ebrahimi Warkiani M 2021 A 3D-printed microfluidic platform for simulating the effects of CPAP on the nasal epithelium Biofabrication 13

[[31] Brewington J J, Filbrandt E T, LaRosa F J, Moncivaiz J D, Ostmann A J, Strecker L M and Clancy J P 2018 Generation of human nasal epithelial cell spheroids for individualized cystic fibrosis transmembrane conductance regulator study Journal of Visualized Experiments 2018

[[32] Bridges M A, Walker D C, Harris R A, Wilson B R and Davidson A G 1991 Cultured human nasal epithelial multicellular spheroids: polar cyst-like model tissues. Biochem Cell Biol 69

[33.] Foty R 2011 A simple hanging drop cell culture protocol for generation of 3D spheroids Journal of Visualized Experiments

[[34] Jørgensen A, Young J, Nielsen J E, Joensen U N, Toft B G, Rajpert-De Meyts E and Loveland K L 2014 Hanging drop cultures of human testis and testis cancer samples: A model used to investigate activin treatment effects in a preserved niche Br J Cancer 110

[[35] Fulcher M L, Gabriel S, Burns K A, Yankaskas J R and Randell S H 2005 Well-differentiated human airway epithelial cell cultures. Methods Mol Med 107

[36.] Srinivasan B, Kolli A R, Esch M B, Abaci H E, Shuler M L and Hickman J J 2015 TEER Measurement Techniques for In Vitro Barrier Model Systems J Lab Autom 20

[37.] Smith C M, Djakow J, Free R C, Djakow P, Lonnen R, Williams G, Pohunek P, Hirst R A, Easton A J, Andrew P W and O’Callaghan C 2012 CiliaFA: A research tool for automated, high-throughput measurement of ciliary beat frequency using freely available software Cilia 1

[38.] Zheng G X Y, Terry J M, Belgrader P, Ryvkin P, Bent Z W, Wilson R, Ziraldo S B, Wheeler T D, McDermott G P, Zhu J, Gregory M T, Shuga J, Montesclaros L, Underwood J G, Masquelier D A, Nishimura S Y, Schnall-Levin M, Wyatt P W, Hindson C M, Bharadwaj R, Wong A, Ness K D, Beppu L W, Deeg H J, McFarland C, Loeb K R, Valente W J, Ericson N G, Stevens E A, Radich J P, Mikkelsen T S, Hindson B J and Bielas J H 2017 Massively parallel digital transcriptional profiling of single cells Nat Commun 8

[39.] Perez J T, Garćia-Sastre A and Manicassamy B 2013 Insertion of a GFP Reporter Gene in Influenza Virus Curr Protoc Microbiol 015

[40.] Schulze-Horsel J, Genzel Y and Reichl U 2008 Flow cytometric monitoring of influenza A virus infection in MDCK cells during vaccine production BMC Biotechnol 8

[41.] Zamora J L R and Aguilar H C 2018 Flow virometry as a tool to study viruses Methods 134–135

[42.] Li W, Germain R N and Gerner M Y 2019 High-dimensional cell-level analysis of tissues with Ce3D multiplex volume imaging Nat Protoc 14

[43.] Tokuda S, Higashi T and Furuse M 2014 ZO-1 knockout by talen-mediated gene targeting in MDCK cells: Involvement of ZO-1 in the regulation of cytoskeleton and cell shape PLoS One 9

[44.] Wang D Y, Li Y, Yan Y, Li C and Shi L 2015 Upper Airway Stem Cells: Understanding the Nose and Role for Future Cell Therapy Curr Allergy Asthma Rep 15

[45.] Baldassi D, Gabold B and Merkel O M 2021 Air−Liquid Interface Cultures of the Healthy and Diseased Human Respiratory Tract: Promises, Challenges, and Future Directions Adv Nanobiomed Res 1

[46.] Garcıá S R, Deprez M, Lebrigand K, Cavard A, Paquet A, Arguel M J, Magnone V, Truchi M, Caballero I, Leroy S, Marquette C H, Marcet B, Barbry P and Zaragosi L E 2019 Novel dynamics of human mucociliary differentiation revealed by single-cell RNA sequencing of nasal epithelial cultures Development (Cambridge) 146

[47.] Diamond G, Legarda D and Ryan L K 2000 The innate immune response of the respiratory epithelium Immunol Rev 173

[48.] Whitsett J A and Alenghat T 2015 Respiratory epithelial cells orchestrate pulmonary innate immunity Nat Immunol 16

[49.] Fishman A P 2008 Fishman’s Pulmonary Diseases and Disorders (4th Ed.) vol 4

[50.] Jaspers I, Ciencewicki J M, Zhang W, Brighton L E, Carson J L, Beck M A and Madden M C 2005 Diesel exhaust enhances influenza virus infections in respiratory epithelial cells Toxicological Sciences 85

[51.] Jaspers I, Flescher E and Chen L C 1997 Ozone-induced IL-8 expression and transcription factor binding in respiratory epithelial cells Am J Physiol Lung Cell Mol Physiol 272

[52.] Kesic M J, Meyer M, Bauer R and Jaspers I 2012 Exposure to ozone modulates human airway protease/antiprotease balance contributing to increased influenza a infection PLoS One 7

[53.] Roth M, Steiner S, Bisig C, Comte P, Czerwinski J, Rothen-Rutishauser B, Latzin P and Müller L 2015 Effect of gasoline exhaust emission on bronchial epithelial cells and natural killer cells

[54.] Horvath K M, Brighton L E, Zhang W, Carson J L and Jaspers I 2011 Epithelial cells from smokers modify dendritic cell responses in the context of influenza infection Am J Respir Cell Mol Biol 45

[55.] Reeves S R, Barrow K A, White M P, Rich L M, Naushab M and Debley J S 2018 Stability of gene expression by primary bronchial epithelial cells over increasing passage number BMC Pulm Med 18

[[56] de Jong P M, van Sterkenburg M A J A, Kempenaar J A, Dijkman J H and Ponec M 1993 Serial culturing of human bronchial epithelial cells derived from biopsies In Vitro Cellular & Developmental Biology – Animal: Journal of the Society for In Vitro Biology 29

[57.] Ehrhardt C, Collnot E M, Baldes C, Becker U, Laue M, Kim K J and Lehr C M 2006 Towards an in vitro model of cystic fibrosis small airway epithelium: Characterisation of the human bronchial epithelial cell line CFBE41o-Cell Tissue Res 323

[[58] Bauer R N, Brighton L E, Mueller L, Xiang Z, Rager J E, Fry R C, Peden D B and Jaspers I 2012 Influenza enhances caspase-1 in bronchial epithelial cells from asthmatic volunteers and is associated with pathogenesis Journal of Allergy and Clinical Immunology 130

[59.] Devalia J L, Sapsford R J, Wells C W, Richman P and Davies R J 1990 Culture and comparison of human bronchial and nasal epithelial cells in vitro Respir Med 84

[60.] Horvath K M, Herbst M, Zhou H, Zhang H, Noah T L and Jaspers I 2011 Nasal lavage natural killer cell function is suppressed in smokers after live attenuated influenza virus Respir Res 12

[[61] Heffler E, Landi M, Caruso C, Fichera S, Gani F, Guida G, Liuzzo M T, Pistorio M P, Pizzimenti S, Riccio A M, Seccia V, Ferrando M, Malvezzi L, Passalacqua G and Gelardi M 2018 Nasal cytology: Methodology with application to clinical practice and research Clinical and Experimental Allergy 48

[62.] Meltzer E O and Jalowayski A A 1988 Nasal Cytology in Clinical Practice Am J Rhinol 2

[[63] Prior A J, Calderon M A Lavelle R J and Davies R J 1995 Nasal biopsy: indications, techniques and complications Respir Med 89

[64.] Becker U, Ehrhardt C, Schneider M, Muys L, Gross D, Eschmann K, Schaefer U F and Lehr C M 2008 A comparative evaluation of corneal epithelial cell cultures for assessing ocular permeability ATLA Alternatives to Laboratory Animals 36

[[65] Hufnagel M, May N, Wall J, Wingert N, Garcia-Käufer M, Arif A, Hübner C, Berger M, Mülhopt S, Baumann W, Weis F, Krebs T, Becker W, Gminski R, Stapf D and Hartwig A 2021 Impact of nanocomposite combustion aerosols on a549 cells and a 3d airway model Nanomaterials 11

[[66] Talikka M, Kostadinova R, Xiang Y, Mathis C, Sewer A, Majeed S, Kuehn D, Frentzel S, Merg C, Geertz M, Martin F, Ivanov N v., Peitsch M C and Hoeng J 2014 The Response of Human Nasal and Bronchial Organotypic Tissue Cultures to Repeated Whole Cigarette Smoke Exposure Int J Toxicol 33

[67.] Rotoli B M, Barilli A, Visigalli R, Ferrari F, Frati C, Lagrasta C A, di Lascia M, Riccardi B, Puccini P and Dall’asta V 2020 Characterization of ABC transporters in epiairway, a cellular model of normal human bronchial epithelium Int J Mol Sci 21

[68.] Matai I, Kaur G, Seyedsalehi A, McClinton A and Laurencin C T 2020 Progress in 3D bioprinting technology for tissue/organ regenerative engineering Biomaterials 226

[69.] Mandrycky C, Wang Z, Kim K and Kim D H 2016 3D bioprinting for engineering complex tissues Biotechnol Adv 34

[70.] Ayan B, Heo D N, Zhang Z, Dey M, Povilianskas A, Drapaca C and Ozbolat I T 2020 Aspiration-assisted bioprinting for precise positioning of biologics Sci Adv 6 eaaw5111

[[71] Zhang J, Wehrle E, Rubert M and Müller R 2021 3d bioprinting of human tissues: Biofabrication, bioinks and bioreactors Int J Mol Sci 22

[72.] Dubbin K, Hori Y, Lewis K K and Heilshorn S C 2016 Dual-Stage Crosslinking of a Gel-Phase Bioink Improves Cell Viability and Homogeneity for 3D Bioprinting Adv Healthc Mater 5

[73.] Pössl A, Hartzke D, Schlupp P and Runkel F E 2022 Optimized cell mixing facilitates the reproducible bioprinting of constructs with high cell viability Applied Sciences (Switzerland) 12

[74.] Kim B S, Lee J S, Gao G and Cho D W 2017 Direct 3D cell-printing of human skin with functional transwell system Biofabrication 9

[[75] Tan K sen, Ong H H, Yan Y, Liu J, Li C, Ong Y K, Thong K T Choi H W, Wang D Y and Chow V T 2018 In Vitro Model of Fully Differentiated Human Nasal Epithelial Cells Infected with Rhinovirus Reveals Epithelium-Initiated Immune Responses Journal of Infectious Diseases 217

[76.] Dimova S, Brewster M E, Noppe M, Jorissen M and Augustijns P 2005 The use of human nasal in vitro cell systems during drug discovery and development Toxicology in Vitro 19

[[77] Aydin M, Naumova E A, Bellm A, Behrendt A K, Giachero F, Bahlmann N, Zhang W, Wirth S, Paulsen F, Arnold W H and Ehrhardt A 2021 From submerged cultures to 3d cell culture models: Evolution of nasal epithelial cells in asthma research and virus infection Viruses 13

[[78] Jafri H S, Chávez-Bueno S, Mejías A, Gómez A M, Ríos A M, Nassi S S, Yusuf M, Kapur P, Hardy R D, Hatfield J, Rogers B B, Krisher K and Ramilo O 2004 Respiratory syncytial virus induces pneumonia, cytokine response, airway obstruction, and chronic inflammatory infiltrates associated with long-term airway hyperresponsiveness in mice Journal of Infectious Diseases 189

[79.] Rogers D F 2003 The airway goblet cell International Journal of Biochemistry and Cell Biology 35

[80.] Groves H E, Guo-Parke H, Broadbent L, Shields M D and Power U F 2018 Characterisation of morphological differences in well-differentiated nasal epithelial cell cultures from preterm and term infants at birth and one-year PLoS One 13

[81.] Kreft M E, Hudoklin S, Jezernik K and Romih R 2010 Formation and maintenance of blood-urine barrier in urothelium Protoplasma 246

[82.] Matter K and Balda M S 2003 Functional analysis of tight junctions Methods 30

[83.] Kreft M E, Tratnjek L, Lasič E, Hevir N, Rižner T L and Kristan K 2020 Different Culture Conditions Affect Drug Transporter Gene Expression, Ultrastructure, and Permeability of Primary Human Nasal Epithelial Cells Pharm Res 37

[[84] Broadbent L, Manzoor S, Zarcone M C, Barabas J, Shields M D, Saglani S, Lloyd C M, Bush A, Custovic A, Ghazal P, Gore M, Marsland B, Roberts G, Schwarze J, Turner S and Power U F 2020 Comparative primary paediatric nasal epithelial cell culture differentiation and RSV-induced cytopathogenesis following culture in two commercial media PLoS One 15

[85.] Jorissen M 1998 Correlations among mucociliary transport, ciliary function, and ciliary structure Am J Rhinol 12

[86.] Wilson R, Sykes D A, Currie D and Cole P J 1986 Beat frequency of cilia from sites of purulent infection Thorax 41

[87.] Nuutinen J, Rauch-Toskala E, Saano V and Joki S 1993 Ciliary Beating Frequency in Chronic Sinusitis Arch Otolaryngol Head Neck Surg 119

[88.] Tratnjek L, Kreft M, Kristan K and Kreft M E 2020 Ciliary beat frequency of in vitro human nasal epithelium measured with the simple high-speed microscopy is applicable for safety studies of nasal drug formulations Toxicology in Vitro 66

[89.] Mallants R, Jorissen M and Augustijns P 2010 Beneficial effect of antibiotics on ciliary beat frequency of human nasal epithelial cells exposed to bacterial toxins Journal of Pharmacy and Pharmacology 60

[90.] Chakraborty J, Banerjee I, Vaishya R and Ghosh S 2020 Bioengineered in Vitro Tissue Models to Study SARS-CoV-2 Pathogenesis and Therapeutic Validation ACS Biomater Sci Eng 6

[91.] Heinen N, Klöhn M, Steinmann E and Pfaender S 2021 In vitro lung models and their application to study sars-cov-2 pathogenesis and disease Viruses 13

[92.] Richard M, van den Brand J M A, Bestebroer T M, Lexmond P, de Meulder D, Fouchier R A M, Lowen A C and Herfst S 2020 Influenza A viruses are transmitted via the air from the nasal respiratory epithelium of ferrets Nat Commun 11

[93.] Tran B M, Grimley S L, McAuley J L, Hachani A, Earnest L, Wong S L, Caly L, Druce J, Purcell D F J, Jackson D C, Catton M, Nowell C J, Leonie L, Deliyannis G, Waters S A, Torresi J and Vincan E 2022 Air-Liquid-Interface Differentiated Human Nose Epithelium: A Robust Primary Tissue Culture Model of SARS-CoV-2 Infection Int J Mol Sci 23

[94.] Tavafoghi M, Khademhosseini A and Ahadian S 2021 Advances and challenges in bioprinting of biological tissues and organs Artif Organs 45

