## Supplementary Information for "High-Throughput Bioprinting of the Nasal Epithelium using Patient-derived Nasal Epithelial Cells"

**Supplementary Tables**

**Table S1.** Reagents and antibodies used in this study.

| **Reagent** | **Source** | **Catalog number** |
| --- | --- | --- |
| **Chemical, media, and other reagents** | | |
| Paraformaldehyde 16% | Thermofischer | 28906 |
| Triton X-100 | Sigma | 9036-19-5 |
| Bovine Serum Albumin | Research Products International Corp. | 1004519 |
| Glycine | Sigma | 207300-76-3 |
| Tween 20 | Sigma | 9005-64-5 |
| Normal Goat Serum | Abcam | ab7481 |
| Phalloidin ATTO647N | Sigma | 65906 |
| Fluromount-G | Thermofischer | 00-4958-02 |
| DAPI | Thermofischer | D1306 |
| Fc receptor blocker | Innovex | NB309 |
| Background buster | Innovex | NB306 |
| **Antibodies** |  |  |
| Anti-SCGB1A1 (rat IgG) | R+D | MAB4218 |
| Anti-ZO-1 Monoclonal | Invitrogen | 33-9100 |
| Alpha Tubulin (TUBA4A) | Origen | TA385483 |
| Anti-acetylated alpha-tubulin (mouse IgG2b) | Invitrogen | 32-2700 |
| Anti-MUC5AC | Invitrogen | MA5-12178 |
| Anti-MUC5AC AF700 | Novus | NBP2-32732AF700 |
| Anti-cytokeratin 5 AF594 | Novus | NBP2-61931AF594 |
| Anti-mouse IgG2b AF555 | Invitrogen | A-21147 |
| Anti-rat IgG AF488 | Invitrogen | A-11006 |
| Anti-mouse IgG AF488 | Invitrogen | A-11017 |

**Table S2.** Selected cell marker genes expressed in bioprinted nasal samples for cluster identification.

| **Cell Type** | **mRNA** |
| --- | --- |
| Basal | DLK2, KRT14, LAMA3, TP63 |
| Suprabasal | KRT5, KRT13 |
| Club | FABP5 |
| Goblet | MUC1, MUC5AC, MUC5B, SCGB1A1 |
| Multiciliated | FOXJ1, IFT57, LRRC23, PIFO |

**Supplementary Figures**

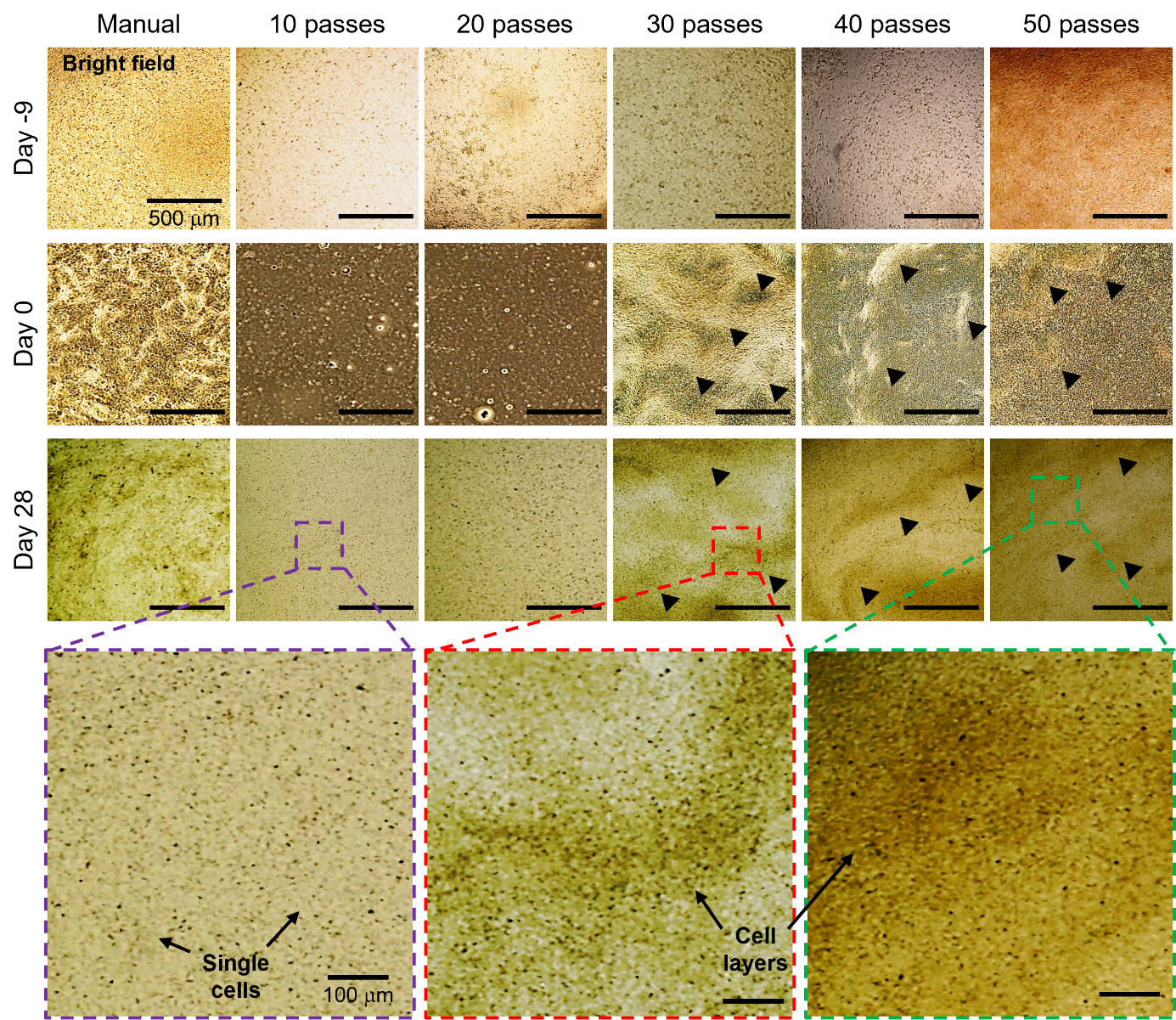

**Figure S1.** Optical images of hNECs processed with manual seeding and DBB using different conditions at Days -9, 0, and 28. Black arrows indicate agglomerated cell layers.

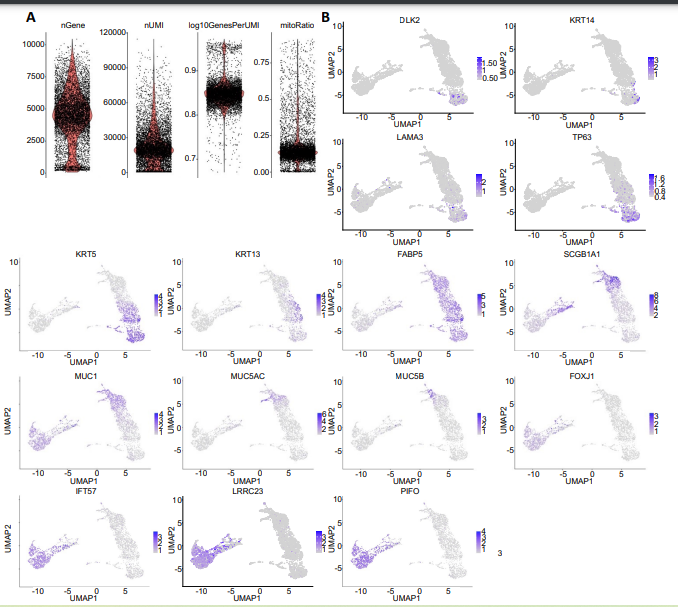

**Figure S2.** scRNA-seq quality controls and differential gene expressions **A)**Violin plots showing genes per cell, unique molecular identifiers (UMI) per cell, number of genes per UMI for each cell and mitochondrial (mito) genes per total cell genes before quality control assessment. **B)** UMAP plot overlays showing selected gene expression distribution across clusters.

**Supplementary Videos**

**Video S1:** High-throughput DBB in 2x speed.

**Video S2:** Ciliary Beating of manual seeding group (1x speed), 30-pass group (1x speed), 40-pass group (1x speed),50-pass group (1x speed and 0.5x speed). All videos were captured with 40x microscope magnification.
